# PI3Kδ inhibition supports memory T cells with enhanced antitumor fitness

**DOI:** 10.1101/166074

**Authors:** Jacob S. Bowers, Kinga Majchrzak, Michelle H. Nelson, Bulent Arman Aksoy, Megan M. Wyatt, Aubrey S. Smith, Stefanie R. Bailey, Lillian Neal, Jeff Hammerbacher, Chrystal M. Paulos

## Abstract

Phosphatidylinositol-3-kinase p110δ (PI3Kδ) inhibition by Idelalisib (CAL-101) in hematological malignancies directly induces apoptosis in cancer cells and disrupts immunological tolerance by depleting regulatory T cells (Tregs). Yet, little is known about the direct impact of PI3Kδ blockade on effector T cells from CAL-101 therapy. Herein, we demonstrate a direct effect of p110δ inactivation via CAL-101 on murine and human CD8^+^ T cells that promotes a strong undifferentiated memory phenotype (elevated CD62L/CCR7, CD127 and Tcf7). These CAL-101 T cells also persisted longer after transfer and exerted stronger antitumor immunity compared to traditionally expanded CD8^+^ T cells in two solid tumor models. Thus, this report describes a novel direct enhancement of CD8^+^ T cell memory by a p110δ inhibitor that leads to markedly improved tumor regression. This finding has significant implications to improve outcomes from next generation cancer immunotherapies.

**Highlights:**

- *In vitro* blockade of PI3K p110δ with CAL-101 endows antitumor T cells with a stronger memory phenotype than those treated with AKTi
- The strong memory phenotype of CAL-101 treated cells translates into improved survival of mice bearing aggressive tumors after adoptive transfer of these T cells
- Human CAR engineered T cells treated with CAL-101 possess an enhanced memory phenotype and robust antitumor efficacy
- The antitumor efficacy of CAL-101 primed T cells is not mediated by high CD62L or CD127 expression, but is likely driven by their stem memory phenotype

**eTOC Blurb:** Bowers et al report a novel function of PI3K blockade using the p110δ subunit inhibitor CAL-101 to induce memory and antitumor potency in CD8^+^ T cells. *Ex vivo* treatment of T cells with CAL-101 leads to improved antitumor control and subject survival in both murine transgenic T cell and human CAR T cell models.

## INTRODUCTION

Adoptive T cell transfer therapy (ACT) for cancer enriches and expands autologous tumor-reactive T cells before returning them to the patient. Thus, ACT allows for the *in vitro* selection or generation of T cells optimally suited to exert antitumor immunity *in vivo*, such as memory T cells. Less differentiated memory T cells (those that express high lymphoid homing molecules CD62L and CCR7), mediate robust antitumor responses resulting in better patient outcomes (Sommermeyer et al., 2016; Wang et al., 2016). Conversely, not only are fully differentiated effector CD8^+^ T cells ineffective at clearing tumors (Gattinoni et al., 2005), but these effector cells will corrupt the antitumor potential of less differentiated T cells if cultured *ex vivo* together (Klebanoff et al., 2016). The improved antitumor efficacy of less differentiated T cells is due in part to an improved capacity to engraft and persist long-term in the host. This T cell longevity may be due not only to improved trafficking to lymphoid tissues, but also improved capacity to respond to homeostatic cytokines (Johnson et al., 2015). Additionally, stem memory T cells (Tscm), which are the least differentiated of the memory subsets and express active Wnt/β-catenin signaling, possess the greatest capacity to clear tumors and provide long-term immunity (Gattinoni et al., 2011).

A major pursuit in the cellular therapy field is to preferentially expand large number of T cells possessing a less differentiated state. Particularly exciting are methods that use small molecules that pharmaceutically enhance T cell memory during *ex vivo* expansion. Multiple emerging strategies include blockade of β-catenin degradation (Gattinoni et al., 2011), increasing/strengthening mitochondrial networks to mimic those seen in memory T cells (Buck et al., 2016), denial of glucose via a small molecule inhibitor (Sukumar et al., 2013), and inhibition of the PI3K/AKT axis (Crompton et al., 2015; van der Waart et al., 2014).

The PI3K/AKT axis plays an integral role in T cell activation downstream of the TCR and co-stimulatory molecules. This pathway is important for T cell clonal expansion, survival, and cytokine production (Soond et al., 2010). The PI3K/Akt axis is also involved in memory formation, as AKT phosphorylates and sequesters FOXO transcription factors preventing transcription of CD62L, CCR7, CD127 and other molecules associated with less differentiated T cells (Hedrick et al., 2012). CD8^+^ T cells from patients who have an overactive p110δ rapidly proliferate and become terminally differentiated, leading to chronic inflammation and greater susceptibility to viral infection (Elkaim et al., 2016; Lucas et al., 2014). Yet this pathway is important for long term memory formation of T cells, as T cells from mice with an inactive mutant p110δ subunit proliferate poorly and are less functional (Gracias et al., 2016; Liu et al., 2009). In particular, the few surviving memory cells in these mice are insufficient to mount a successful response to reinfection (Liu and Uzonna, 2010; Pearce et al., 2015).

Even though the PI3K/AKT pathway plays such a central role in effector T cell biology, the impact of p110δ inhibitors on cancer immunity has historically been associated solely with the promotion of effector CD8^+^ T cells due to ablating Treg suppression (Ali et al., 2014). Yet, recent studies now clearly show that adoptively transferred T cells treated with a small molecule inhibitor of AKT, AKT inhibitor VIII (AKTi), exert stronger antitumor responses in both GVL and melanoma (Crompton et al., 2015; van der Waart et al., 2014). Moreover, we reported that PI3Kδ inhibition with CAL-101 in a Th17 culture manifests a precursor lymphocyte population with a central memory-like phenotype and enhanced antitumor activity (Majchrzak et al., 2017). It remains unknown if the improved treatment outcome by this CAL-101 therapy is due to the selective Treg depletion via CAL-101 or if it directly regulates effector CD8^+^ T cell memory. Collectively, both the genetic and pharmaceutical studies indicate a direct role of PI3K blockade on effector CD8^+^ T cells memory. Yet, while genetic perturbations seem to reduce effector memory capacity, pharmaceutical inhibition appears to enhance memory. Additionally, though PI3K and AKT inhibition are often considered identical because of their linear signaling relationship, PI3K inhibition may change effector T cell physiology differently than AKT inhibition. For example, PI3K interacts with multiple other kinases besides AKT including MAPK and PKC kinases (Hoeflich et al., 2009; Le Good et al., 1998; Yu et al., 2002), as well as other AGC family kinases through the downstream kinase PDK1 (Pearce et al., 2010). We therefore posited that pharmaceutical inhibition of p110δ might directly endow tumor-reactive CD8^+^ T cells with more durable memory properties than direct AKT inhibition.

We tested our hypothesis in both murine transgenic TCR pmel-1 CD8^+^ T cells and human peripheral blood T cells engineered with a tumor antigen specific chimeric antigen receptor (CAR). These cells were expanded with either the P110δ inhibitor CAL-101 or AKTi before adoptive transfer. We found that CAL-101 expanded central memory T cells (both murine and human) with even greater levels of CCR7, CD62L, and the alpha receptor for IL-7 (CD127) than AKTi. CAL-101 treated T cells had the highest levels of donor cells post-transfer, which corresponded with improved tumor control and lengthened survival. While CAL-101 treatment enriched T cells with high lymphoid homing receptors and responsiveness to IL-7, this could not explain the improved antitumor efficacy. CD62L^+^ T cells sorted from IL-2 expanded T cells could not recapitulate the potency of CAL-101 priming or naïve T cells. Additionally, depletion of endogenous IL-7 in the host did not abrogate the capacity of CAL-101 primed T cells to clear tumor. However, deep sequencing of mRNA indicated that PI3Kδ blockade uniquely upregulated the stem memory transcription factor Tcf7 much more than AKT inhibition. These results imply that the simple application of CAL-101 in the preparation of effector T cells directly induces a memory program that greatly improves their capacity to control malignancies.

## RESULTS

### Pharmaceutical inhibition of p110δ in pmel-1 CD8^+^ T cells increases both CD62L and CD127 expression without hindering expansion

To test if PI3Kδ inhibition enhanced the antitumor capacity of T cells similarly to direct AKT inhibition, CD8^+^ T cells from pmel-1 transgenic mice (CD8^+^ T cells with a transgenic TCR specific for the melanoma/melanocyte antigen gp100) were activated with their cognate antigen and treated with CAL-101 throughout culture. As controls, T cells were expanded without drug (vehicle) or with AKT inhibitor VIII (AKT 1/2 hereafter denoted as AKTi), a published method to enhance T cell memory and antitumor efficacy (Crompton et al., 2015). As expected, drug treatment induced marked differences in memory. While 65% of vehicle treated CD8^+^ T cells expressed CD62L, nearly all cells treated with AKTi or with CAL-101 expressed CD62L (Figure 1A). However, CAL-101 exerted a stronger impact on the cells than direct AKT inhibition. The MFI of CD62L on CAL-101 treated T cells was higher than on those treated with AKTi. While both drugs decreased the activation marker CD69, only CAL-101 reduced CD44 expression (Figure 1B). While this CD44^lo^CD62L^hi^ phenotype mirrored the phenotype of a naïve T cells, CAL-101 treated cells expanded similarly to both vehicle and AKTi treated cells. These data suggest that despite PI3Kδ inhibition, T cell receptor and downstream proliferative signaling were intact in pmel-1 CD8^+^ T cells *in vitro* (Figure 1C).

**Figure 1.**
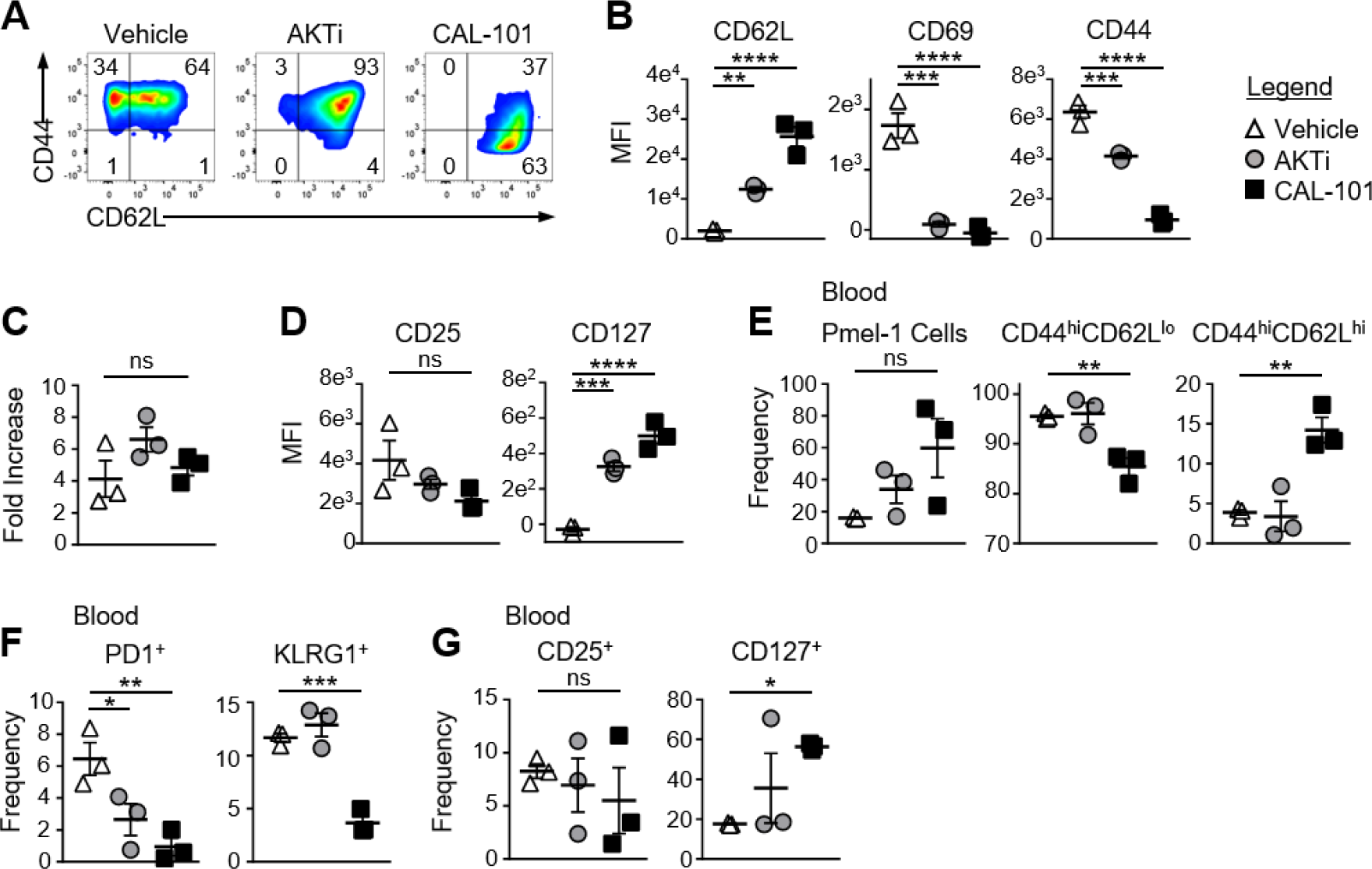
CAL-101 priming supports a de-differentiated memory phenotype including high IL-7Ra expression on pmel-1 CD8^+^ T cells. (A) Representative flow plots of CD44 by CD62L expression on pmel-1 CD8^+^ T cells primed with vehicle, AKTi or CAL-101 after 5 days of culture; representative of 3 independent cultures. (B) MFI of CD62L, CD69 and CD44 on pmel-1 CD8^+^ T cells after 5 days of culture; n=3 independent cultures. (C) Fold increase of pmel-1 CD8^+^ T cells primed with vehicle (DMSO), AKTi, or CAL-101 5 days following antigen stimulation; n=3 independent cultures. (D) MFI of CD25 and CD127 on pmel-1 CD8^+^ T cells after 5 days of culture; n=3 independent cultures. (E-G) Frequency of donor pmel-1 T cells and extracellular markers on pmel-1 donor T cells in blood; n=3 mice/group. One way repeated measures ANOVA; ns = not statistically significant, * = p<0.05, ** = p<0.01, *** = p<0.001, **** = p<0.0001.

### CAL-101 treated CD8^+^ T cells robustly engraft *in vivo* and maintain CD62L and CD127 expression

While AKTi and CAL-101 slightly (but not significantly) reduced CD25 (IL-2 receptor alpha) on pmel-1 CD8^+^ T cells, these small molecules substantially elevated CD127, the alpha receptor for IL-7 (Figure 1D). Since IL-7 signaling is important for naïve and central memory T cell homeostasis (Schluns et al., 2000), and supports donor T cells in the host (Johnson et al., 2015), we posited that the memory T cells generated from either CAL-101 or AKTi treatment would show superior engraftment and subsequent antitumor immunity compared to vehicle. Interestingly, CAL-101 pmel-1 CD8^+^ T cells engrafted with increased frequency in the blood compared to AKTi pmel-1 CD8^+^ T cells (Figure 1E). Additionally, while both AKTi and CAL-101 increased CD62L expression *in vitro* (Figure 1B), only CAL-101 treated pmel-1 retained significant levels of CD44^hi^CD62L^hi^ with fewer CD44^hi^CD62L^lo^ circulating cells (Figure 1E). Although both AKTi and CAL-101 T cells expressed less PD1 than vehicle *in vivo*, only CAL-101 T cells maintained reduced frequencies of the exhaustion marker KLRG1 (Figure 1F). Infused CD8^+^ T cells also retained far more CD127 on their cell surface *in vivo* when primed *ex vivo* with CAL-101 compared to untreated or Akti treated pmel-1 CD8^+^ T cells (Figure 1G). CAL-101 donor T cells were also detected at higher levels within the spleen and draining (inguinal) lymph nodes of tumor-bearing mice (not shown). Thus, we found that CAL-101 treated CD8^+^ T cells robustly engraft *in vivo*, maintain CD62L and CD127 expression and resist exhaustion.

### CAL-101 T cells impair tumor growth and prolong survival

We suspected that the less differentiated memory phenotype of pmel-1 T cells fostered by CAL-101 treatment would translate to their improved ability to infiltrate and regress tumor once infused into melanoma-bearing mice. As expected, CAL-101 treated T cells were detected at higher levels in the tumor compared to control or AKTi treated T cells (Figure 2A). More CAL-101 donor cells expressed a central memory phenotype within the tumor, however, the percentage of PD1^+^ and KLRG1^+^ donor cells was similar between groups (Figure 2A). Importantly, CAL-101 primed T cells were the most effective at slowing growth of melanoma in mice (Figure 2B-C), extending the lifespan of the animals beyond the survival of the vehicle or AKTi groups (Figure 2D). In fact, the tumor control exerted by AKTi treated T cells was only slightly improved over tumor control by vehicle treated T cells (Figure 2C). Since we were using a 10-fold higher drug dose in the CAL-101 cultures compared to the published amount of AKTi (10μM vs 1μM) we posited that the difference in antitumor activity could be due to higher AKT inhibition by the elevated concentration of CAL-101. To test this idea, we treated pmel-1 CD8 T cells with either 1μM or 10μM of AKTi or CAL-101. Increasing the amount of AKTi to 10μM marginally improved the antitumor efficacy of the T cells similar to treatment with 1μM CAL-101 (Figure S1-A). However, 10μM CAL-101 treatment markedly improved tumor control and significantly improved survival compared to both 10μM AKTi and 1μM CAL-101 (Figure S1A-B). Additionally, while neither 10μM AKTi or 10μM CAL-101 impaired the logarithmic expansion of mouse pmel-1 CD8^+^ T cells (not shown), we found that 10μM AKTi dramatically inhibited growth of human T cells (Figure S1C). Thus, as high dose AKTi did not profoundly improve antitumor T cell potency against large melanoma tumors but did compromise the overall yield of human T cell cultures, we proceeded with comparisons of 1μM AKTi to 10μM CAL-101 in our remaining studies.

**Figure 2.**
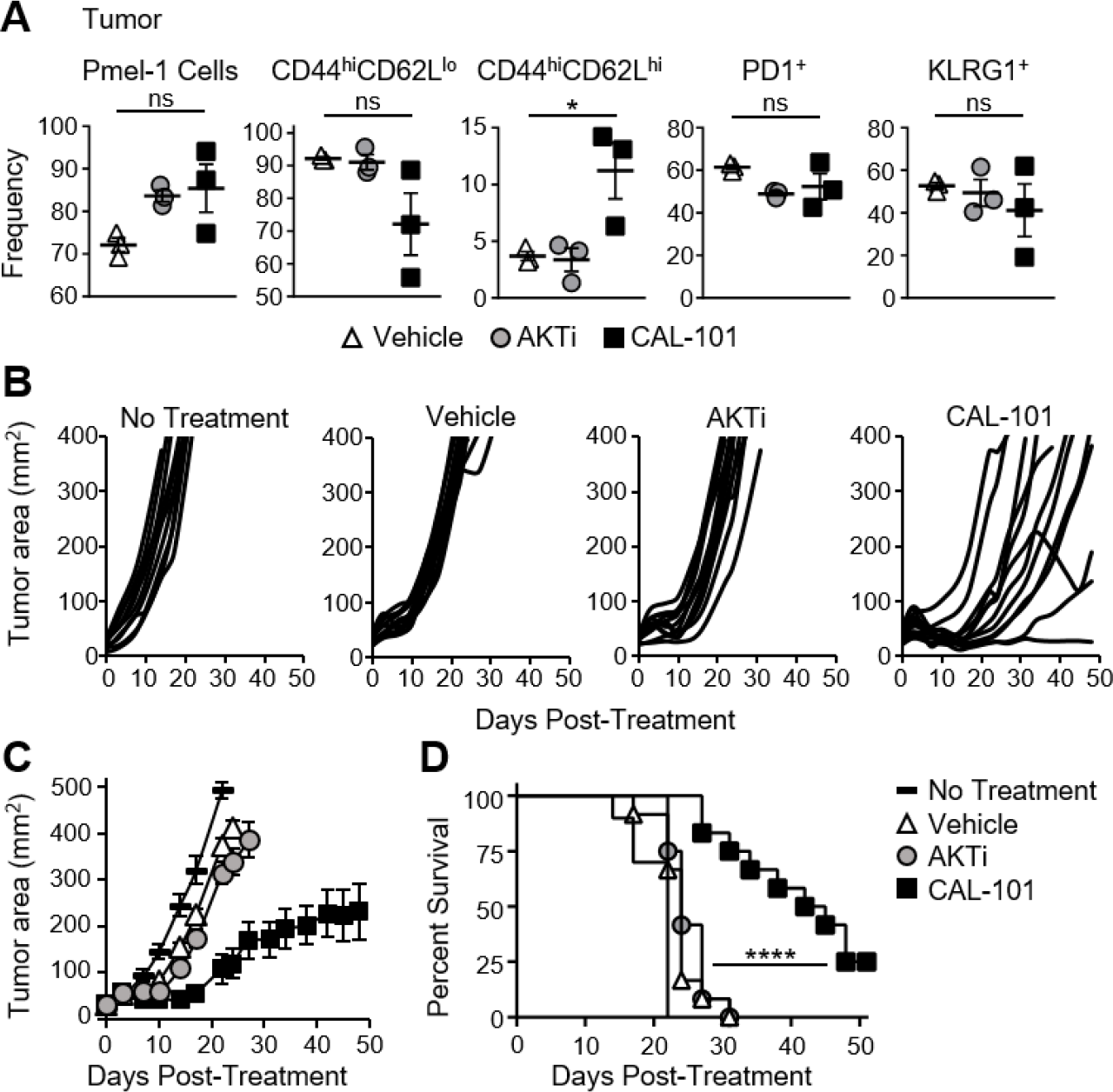
CAL-101 primed pmel-1 CD8^+^ T cells exert stronger antitumor response against B16F10 tumors. (A) Frequency of donor pmel-1 T cells and extracellular markers on pmel-1 (vβ13^+^) donor T cells in tumor 7 days following treatment; n=3 mice/group. One way repeated measures ANOVA; ns = not statistically significant, * = p<0.05. Tumor burden (mm^2^) of (B) individual mice and (C) Average tumor burden (mm^2^) of treatment groups which received no T cell treatment, or 8 x 10^5^ pmel-1 CD8^+^ T cells primed with vehicle, AKTi, or CAL-101 *ex vivo*; n=10-12 mice/group. (D) Percent survival of above treatment groups. Kaplan Meier curve analyzed by log rank test; **** = p<0.0001. (See also Figure S1).

### CAL-101 induces a stronger central memory phenotype than AKTi

We next examined if inhibiting PI3Kδ would augment the fitness and memory properties of human T cells. To do this, human CD3^+^ T cells were expanded with CD3/CD28 beads under CAL-101 or AKTi and compared to a vehicle control. All three T cell groups expressed similar CD45RO levels, a marker of T cell maturation. Both AKTi and CAL-101 treated T cells had slightly increased CD62L over vehicle, though only CAL-101 primed T cells had significantly higher CCR7 (Figure 3A-B). PD1 was nominally expressed on all groups (not shown), but both drug treatments prevented the up-regulation of co-inhibitory receptor TIM3 post-activation compared to untreated T cells. Interestingly, CAL-101 significantly reduced TIM3 (Figure 3C). PI3Kδ and AKT blockade did not impact the expression of CD25, but only CAL-101 greatly increased CD127 on human T cells (Figure 3C-D). Collectively, our data reveal that CAL-101 supports the generation of human T cells with a central memory phenotype with reduced markers of inhibition compared to treatment with AKTi.

**Figure 3.**
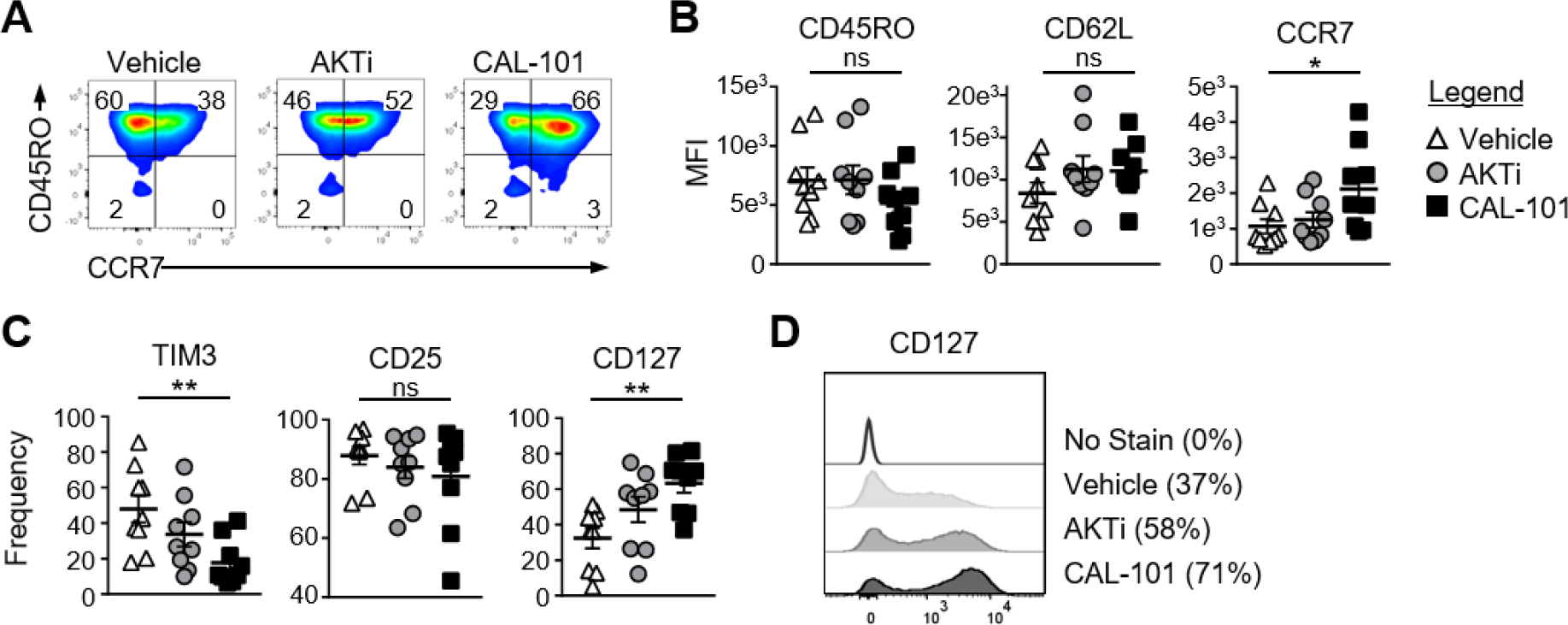
PI3Kδ and AKT blockade induce a central memory phenotype in human CAR CD3^+^ T cells. (A) CD44 and CD62L expression on vehicle, AKTi, or CAL-101 treated T cells from normal donor PBMC; representative of 9 donors. (B) MFI of memory markers and (C) frequency of human CD3^+^ T cells expressing TIM3, CD25 and CD127; n=9 normal donors. One way repeated measures ANOVA; ns = not significant, * = p<0.05, ** = p<0.01. (D) Histogram of CD127 expression on vehicle, AKTi, or CAL-101 treated T cells compared to no stain with frequency of positive cells indicated next to legend; representative of 9 donors.

### PI3Kδ blockade augments the antitumor activity of human CAR-T cells

We posited that human tumor-reactive T cells treated with CAL-101 *in vitro* would control the growth of human tumors in NSG mice better than donor vehicle or AKTi T cells. To address this idea, we transduced human T cells with a lentiviral vector containing a chimeric antigen receptor (CAR) that recognizes mesothelin plus 4-1BB and CD3ξ signaling domains (Carpenito et al., 2009). These CAR T cells were expanded for seven days with CD3/CD28 beads and IL-2 in the presence or absence of CAL-101 or AKTi before transfer into mice bearing subcutaneous M108 mesothelioma tumor. While all treatment groups initially reduced tumor burden in the mice, CAL-101 treated CAR+ T cells exerted longer tumor control compared to the vehice or AKTi treated groups (Figure 4A). Conversely, half of tumors in vehicle T cell treated mice, and a quarter of AKTi T cell treated mice relapsed above 150mm^2^ (Figure 4A). The superior antitumor immunity from CAL-101 primed T cells was further evidenced by the majority of tumors in CAL-101 mice having the smallest mean tumor weight (Figure 4B) and remaining below baseline measurement at the end of study (Figure 4C). Additionally, CAL-101 T cells persisted at significant levels in circulation 55 days after transfer in most treated mice (Figure 4D). This finding with CAL-101 was in stark contrast to vehicle and AKTi T cell treated mice, which showed low levels of persisting cells in the mice (Figure 4D). Thus, priming T cells with CAL-101 improves engraftment, persistence and tumor destruction by human CAR T cells in solid tumors.

**Figure 4.**
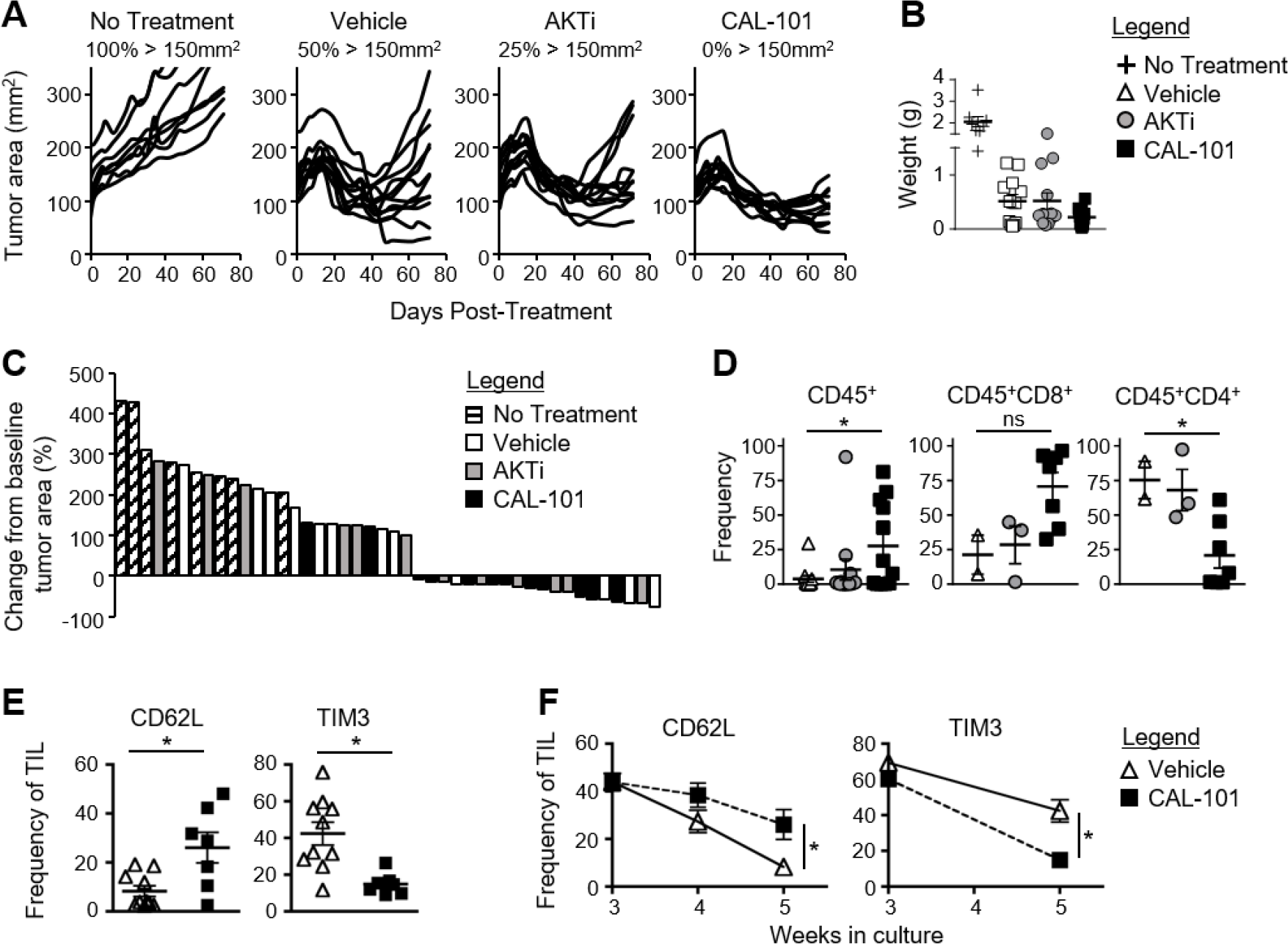
CAL-101 treatment improves tumor control by human CAR T cells compared to vehicle and AKTi treatment. (A) Individual tumor curves of NSG mice given M108 subcutaneously 51 days prior were treated with 4 x 10^5^ CD3^+^ mesoCAR-T cells primed with vehicle, AKTi, or CAL-101; n=8-12 mice/group. (B) Tumor weight at day 71 post-transfer. (C) Percent change in size of tumors 71 days post-treatment compared to baseline tumor measurement at time of treatment. (D) Frequency of human CD45^+^ lymphocytes within the blood of treated mice 55 days post-transfer; n=2-12 mice/group. One way repeated measures ANOVA; ns = not statistically significant, * = p<0.05. (E) Frequency of CD62L^+^ or TIM3^+^ CD3^+^ TIL from lung carcinoma after 5 weeks of growth with either two weeks of CAL-101 treatment or not, and (F) change in frequencies of CD62L^+^ or TIM3^+^ CD3^+^ TIL during drug treatment with CAL-101 or without (vehicle) from week 3 to week 5 of *ex vivo* culture. Groups compared by student’s t-test, * = p<0.05.

Since PI3Kδ inhibition improved the antitumor efficacy of healthy donor derived CAR T cells, we posited that CAL-101 treatment would also improve the memory phenotype of tumor infiltrating lymphocytes (TIL) from patients with lung carcinoma. To address this concept, individual TIL cultures were expanded under IL-2 for three weeks then split into CAL-101 or vehicle treatment groups for another two weeks. We found that CAL-10 supported the generation of TIL with higher CD62L but low TIM3 expression than vehicle after 5 weeks of expansion (Figure 4E). This phenotype appeared to be due to preservation of a less differentiated memory phenotype as vehicle TIL lost expression of CD62L faster than CAL-101 TIL (Figure 4F). Additionally, while TIM3 expression on both groups diminished during *ex vivo* expansion, CAL-101 treatment mediated a rapid loss of TIM3 on TILs (Figure 4F). Collectively, our data reveals that PI3Kδ blockade augments the antitumor activity of human CAR-T cells and fosters the generation of TILs with a less exhausted and preserved memory profile.

### Antitumor potency induced by PI3Kδ inhibition is not due to CD62L

CD62L expression on T cells correlates with improved antitumor immunity in pre-clinical ACT tumor models (Berger et al., 2008; Gattinoni et al., 2005; Hinrichs et al., 2011; Klebanoff et al., 2016; Sommermeyer et al., 2016). Moreover, enriching central memory T cells from peripheral blood and redirecting them with a CD19 specific CAR has shown efficacy in a clinical trial (Wang et al., 2016). Our studies corroborated these reports by showing a correlation between retained CD62L expression *in vivo* by CAL-101 treated donor cells and prolonged tumor control (Figure 1 & 2). Consequently, we naturally hypothesized that CAL-101 induced CD62L on T cells was responsible for their enhanced antitumor potency. Thus, if this concept were true, we suspect that simply sorting the CD62L^+^ T cells from vehicle pmel-1 should mirror the antitumor activity of CAL-101 primed T cells, which express high CD62L. To test this, the following pmel-1 T cell cohorts were administered to B16F10 mice: 1) bulk vehicle T cells (which were 37% CD62L^+^), 2) sorted CD62L^+^ T cells from vehicle treated T cells (98% CD62L+), 3) naïve pmel-1 T cells sorted directly from the spleen (majority CD44^-^CD62L^+^), and 4) CAL-101 treated T cells (which were 97% CD62L^+^, see Figure 5A). Surprisingly, and in contrast to our hypothesis, we found that sorting CD62L^+^ pmel-1 cells from vehicle cultures did not improve treatment outcome over therapy with bulk vehicle. In contrast, CAL-101 mediated prolonged antitumor control (Figure 5B) and better survival (Figure 5C) in mice. Interestingly, naïve sorted T cells closely matched the CAL-101 treated T cells in antitumor potency. It is important to note that despite similar antitumor efficacy, the benefit of CAL-101 expanded T cells over enriched naïve T cells is the potential for propagating signification more cells (i.e. higher cell yield) as vast numbers of cellular product are paramount for successful ACT therapy (Bowers et al., 2017; Gattinoni et al., 2005). Nonetheless, these results imply that PI3Kδ blockade during *in vitro* proliferation might preserve naïve T cell qualities, which would otherwise be corrupted by the expansion process. This finding also corroborates our data from TIL indicating a slower decline in CD62L in CAL-101 treated TIL. Thus, while T cells capable of long-lived memory responses against tumor express CD62L, enriching cells after *ex vivo* expansion that expresses this molecule is not sufficient to drive a successful antitumor response.

**Figure 5.**
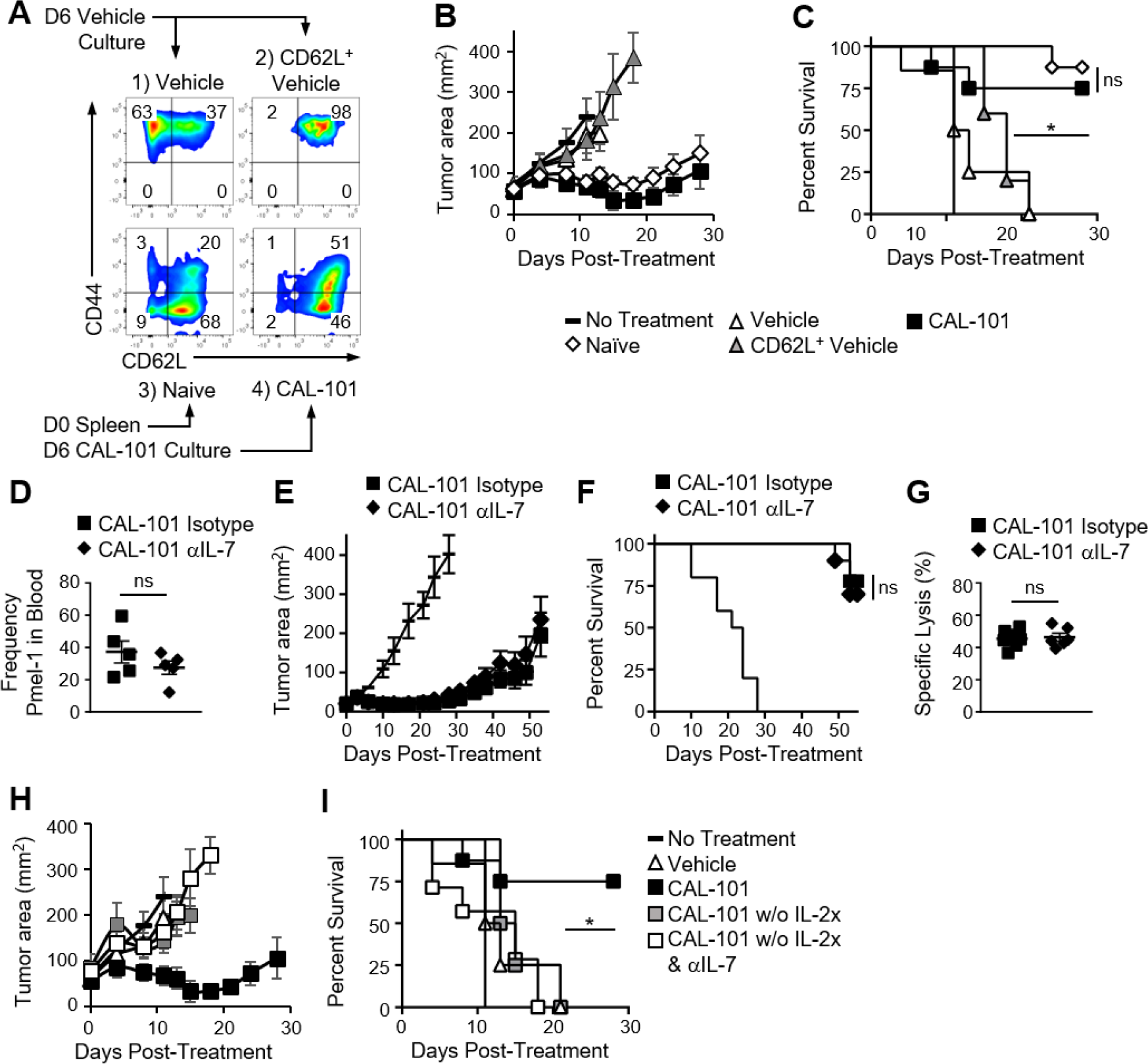
The antitumor potency of CAL-101 primed T cells is CD62L and CD127 independent. (A) Sort diagram with post-sort analysis of CD62L and CD44 expression on pmel-1 T cells. (B) Average tumor burden (mm^2^) and (C) percent survival of mice with B16F10 were treated with 1x10^6^ bulk vehicle, CD62L^+^ vehicle, naïve or CAL-101 pmel-1 cells compared to no treatment; n=5-10 mice/group. Kaplan Meier curve analyzed by log rank test; * = p<0.05. (D) Frequency of donor CAL-101 T cells in mice receiving isotype or IL-7 depleting antibody; n=5 mice/group. One way repeated measures ANOVA; ns = not statistically significant. (See also Figure S2). (E) Average tumor burden and (F) percent survival of of isotype or IL-7 depleted mice receiving CAL-101 primed donor cells; n=10 mice/group. Kaplan Meier curve analyzed by log rank test; ns = not statistically significant. (G) Percent specific lysis of hgp100 loaded splenocytes by CAL-101 primed pmel-1 T cells in isotype treated or IL-7 depleted mice 53 days post-transfer; n=6-7 mice/group. Comparison by student’s t-test; ns = not statistically significant. (H) Average tumor burden and (I) percent survival of mice receiving CAL-101 treated donor cells with IL-2 complex, without IL-2 complex plus isotype, or without IL-2 complex plus IL-7 depletion compared to no treatment; n=7-9 mice/group. Kaplan Meier curve analyzed by log rank test; * = p<0.05.

### CAL-101 primed T cells regress tumor independent of IL-7 signaling

Next we tested if the preserved CD127 on CAL-101 treated T cells was responsible for the robust antitumor properties *in vivo*. PI3Kδ blockade induced CD127 on murine and human T cells *in vitro*. Moreover, CD127 was sustained on CAL-101 treated T cells after transfer (Figure 1E, 2A). We posited that CAL-101 T cells thrived *in vivo* due to their enhanced responsiveness to IL-7. To test the dependence of CAL-101 antitumor responses on IL-7 signaling, we depleted IL-7 in the pmel-1 B16F10 model for the first two weeks after transfer as published (Johnson et al., 2015). We suspected that depleting IL-7 would decrease engraftment of the donor cells and impair their control of tumor growth. In contrast, we found that while CAL-101 primed T cells engrafted at significantly higher numbers than vehicle or AKTi treated T cells in both isotype and IL-7 depleted mice (Figure S2A), there was no significant difference between the CAL-101 treatment groups in frequency or memory phenotype (Figure 5D & S2B-C). CAL-101 primed T cells also exerted their prolonged antitumor immunity whether IL-7 had been depleted or not (Figure 5E) resulting in no differences in survival between the CAL-101 T cell groups (Figure 5F). Furthermore, while IL-7 is important for maintaining naïve and central memory T cell populations, we found no reduction in memory capacity of donor CAL-101 T cells in either isotype or IL-7 depleted animals, as both groups were equally capable of lysing tumor in mice (Figure 5E-F) and ablating hgp100 antigen bearing splenocytes in our very sensitive *in vivo* cytotoxicity assay (Figure 5G).

As IL-2 complex was administered to our melanoma tumor-bearing mice to support the infused CAL-101 T cells, we suspected that this cytokine was important for the engraftment of these infused cells and could compensate for IL-7. IL-2 complex has been reported to support the engraftment and proliferation of CD8^+^ T cells in ACT murine models (Boyman et al., 2006). Thus, we posited that removal of IL-2 complex from the treatment protocol would reveal the importance of IL-7 signaling in the antitumor efficacy mediated by CAL-101 T cells. As expected, tumors grew more rapidly in mice that received CAL-101 T cells without IL-2 complex, compared to those which received both the pmel-1 CD8^+^ T cells and IL-2 complex (Figure 5H). Unexpectedly, however, IL-7 depletion did not compromise the antitumor activity of CAL-101 pmel-1 T cells, regardless of if the mice were treated with IL-2 complex (Figure 5H-I). Collectively, our findings lend further evidence that improved antitumor T cell efficacy from CAL-101 treatment is not due to their increased responsiveness to IL-7 signaling due to high CD127 expression nor was their improved potency due to their heighted expression of CD62L.

### CAL-101 induces stem memory pathways in T cells while AKTi does not

To define how CAL-101 instills infused CD8^+^ T cells with enhanced antitumor activity in vivo, we next sought to use RNA sequencing to uncover the factors potentially responsible for the efficacy of this ACT therapy. We surveyed the differential expression of RNA transcripts of interest associated with memory and effector phenotypes, as well as other pathways influenced by drugs that block PI3K or AKT, including signaling intermediates, metabolic, anti-apoptotic pathways and cell cycle proteins. Similar to our protein data, we found that PI3Kδ inhibition via CAL-101 promoted the up-regulation of multiple central memory markers on T cells, such as CD62L (*Sell*) and CCR7 compared to AKTi or vehicle-treated T cells. CAL-101 also uniquely induced high CD127 (*IL7r*) transcript and stem memory associated transcirpts Lef1 and Tcf7, which were markedly increased compared to AKTi cells (Figure 6A). We expected that a durable memory phenotype would equate to decreased expression of transcripts associated with differentiate T cell effector function. In most cases, as expected, both drug treatments down-regulated effector transcripts including *Fos*, *JunB*, Granzyme B (*Gzmb*) and IFN-γ (*Ifng*), but interestingly the effector transcription factors *Tbx21*, *Eomes* and *Nfatc4* increased with CAL-101 treatment (Figure 6A). We were intrigued by the high expression of stem memory genes *Lef1* and *Tcf7* in CAL-101 treated T cells, known to enhance their persistence and antitumor activity (Bowers et al., 2017; Gattinoni et al., 2011; Majchrzak et al., 2017). We therefore followed up our RNA-seq data by assaying the protein levels of nuclear Lef1, Tcf7 and their upstream molecule β-catenin in CAL-101 treated T cells versus AKTi and vehicle T cells. Similar to our transcript expression results, nuclear β-catenin was similar between groups and both drug treatments had slightly higher, though not statistically significant Lef1. However, PI3Kδ blockade significantly increased nuclear Tcf7 over vehicle while AKTi did not (Figure 6B-C). Thus, our protein results confirm our findings with RNA-seq, and collectively suggest that while the phenotype of AKT inhibited T cells resembles that of potent central memory T cells, PI3Kδ blockade profoundly induced key stem memory transcripts and protein in human CAR T cells that might be fundamentally responsible for their longevity and potency *in vivo*.

**Figure 6.**
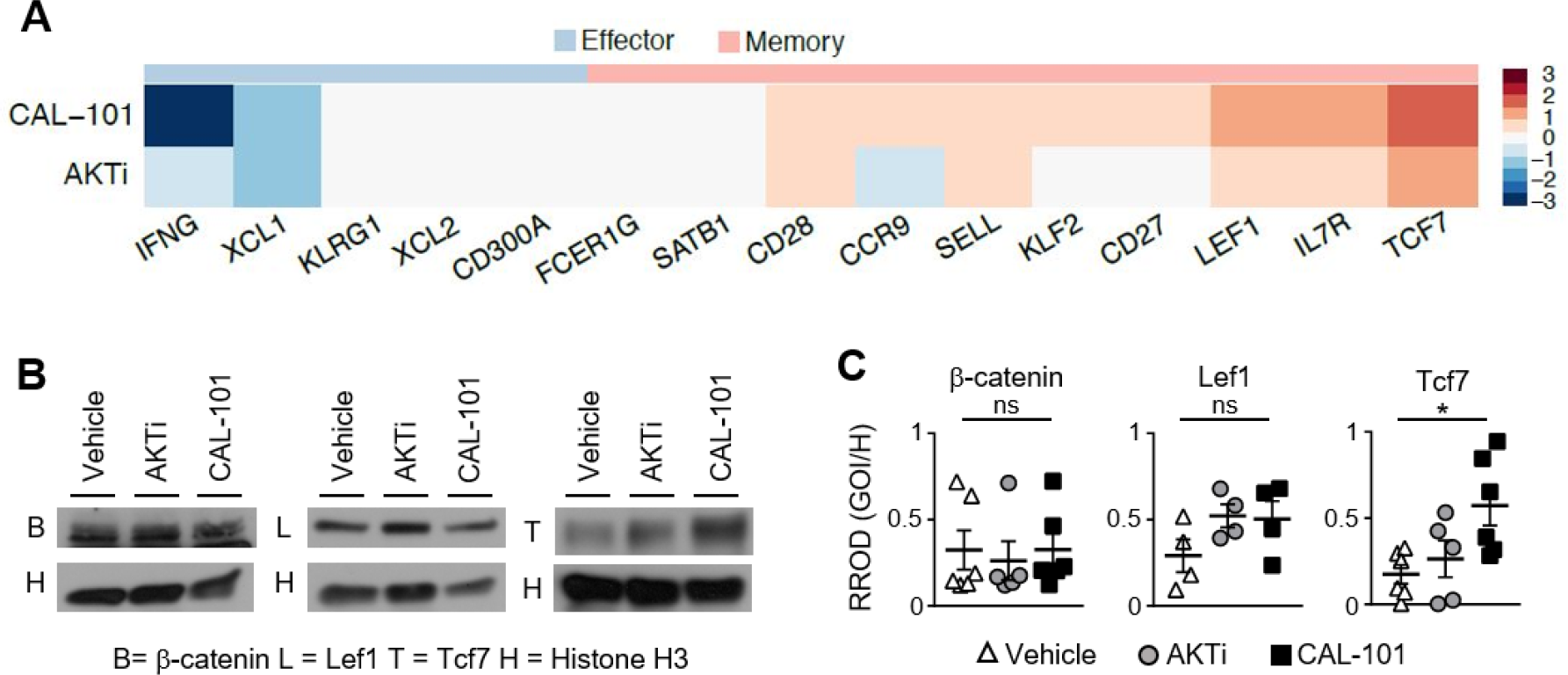
CAL-101 T cells share transcriptional characteristics with stem memory T cells. (A) Differential expression of memory and effector associated genes in CAL-101 or AKTi treated T cells compared to vehicle analyzed using RNA sequencing (See also Figure S3); n=3 normal donors. (B) Western blot of nuclear protein extracts from vehicle, AKTi, or CAL-101 treated T cells, B = b-catenin, L = Lef1, T = Tcf7, H = Histone H3; representative of 4-6 donors. (C) Quantified protein levels relative to Histone H3, RROD GOI/H = relative ratio of optical density (gene of interest over Histone H3); n=4 -6 normal donors. One way repeated measures ANOVA; ns = not statistically significant, * = p<0.05.

In addition to memory transcripts that we hypothesized would be altered by CAL-101 treatment (designated as gene of interest see Figure S3A), we also detected other transcripts which were differentially expressed in CAL-101 primed T cells vs vehicle (designated as differential expression see Figure S3A). many of these genes are associated with T cell fitness versus exhaustion including high anti-apoptotic and metabolism transcripts *Bcl2, stradb* (or ILPIP) and *Ldlrap1* (Figure S3A). Interestingly, ILPIP is an anti-apoptotic protein with energy-generating metabolism (Sanna et al., 2002). Of additional interest, KLF4 was dramatically down regulated in T cell treated with CAL-101. This finding is important as this transcript has been previously reported to restrict memory CD8^+^ T cell responses to foreign antigen (Mamonkin et al., 2013). Collectively based on our data, we suspect that CAL-101 induced Tcf7 signaling while concomitantly down regulating KLF4 to profoundly augment memory and antitumor efficacy by preventing the cells from undergoing terminal differentiation. Thus, CAL-101 regulates distinct pathways (such as Tcf7, KLF4 & ILPIP) potentially critical for supporting durable memory T cell responses to tumors.

## DISCUSSION

Collectively, our data indicate that while PI3K is upstream of AKT, its blockade through inhibition of p110δ induces stronger antitumor capacity in infused CD8^+^ T cells than AKT inhibition. Furthermore, while inhibition of either PI3Kδ or AKT supports the generation of central memory T cells, we found that PI3Kδ blockade preferentially increases CD127 and the stem memory transcription factor Tcf7 while decreasing KLF4. Additionally, PI3Kδ blockade improved the antitumor response of both murine and human tumor-reactive T cells over that of traditionally expanded (vehicle) or AKTi-treated T cells.

Successful immunity against both viral infections and cancer requires long-term protection provided by memory cells (Sallusto et al., 2010). We have classically associated T cell memory with expression of surface markers such as CD62L, CCR7 and CD127 (Sallusto et al., 1999; Schluns et al., 2000), which were dramatically increased in CAL-101 treated T cells. We sought to determine if CAL-101 induced CD62L and CD127 were responsible for the enhanced antitumor immunity after PI3Kδ blockade. We found early evidence to conclude that the longevity of immune responses exerted by CAL-101 T cells cannot be solely explained by high CD62L expression, as sorted CD62L^+^ T cells from vehicle expanded T cells cannot recapitulate the antitumor efficacy of either CAL-101 primed T cells or naïve T cells enriched directly from the spleen. We suspect that uninhibited TCR and IL-2 signals drive vehicle CD62L^+^ cells to rapidly differentiate once transferred *in vivo* (Verdeil et al., 2006). While naïve and CAL-101 T cells exerted similar antitumor immunity, it is important to note that the benefit of CAL-101 treatment is the preservation of a less differentiated phenotype while the T cells proliferate. Expanding tumor-reactive T cells to a high T cell yield for therapy is still vital to the therapy’s success (Bowers et al., 2017; Gattinoni et al., 2005), and thus the low yield of unexpanded naïve cells may be a disadvantage.

Additionally, *in vivo* blockade of IL-7 was insufficient to impair the antitumor efficacy of CAL-101 primed T cells. However, it is probable that the administration of the anti-IL-7 antibody did not completely deplete IL-7 and that the higher expression of CD127 on the CAL-101 primed T cells allowed for successful scavenging of this cytokine. Future studies using T cells from *Il7r* conditional knockout mice may determine the importance of CD127 to the longevity of CAL-101 primed T cell antitumor responses. In contrast, we found that IL-2 complex therapy was crucial to the efficacy of CAL-101 T cells to control tumor growth.

Since PI3Kδ blockade augmented antitumor potency in an apparent CD62L and IL-7 independent manner, we suspected that other aspects of T cell memory and differentiation were being altered uniquely by CAL-101 treatment. One pathway which is consistently associated with durable memory T cells is the Wnt/β-catenin pathway, whether it be active in stem memory T cells (Gattinoni et al., 2011; Gattinoni et al., 2009) or in Th17 cells (Muranski et al., 2011). We found that CAL-101 increased Tcf7 expression in human T cells significantly above vehicle while AKT inhibition did not. Thus, future studies with Tcf7 knockout T cells will help us understand the role and mechanism of Tcf7 in enhancing memory due to PI3Kδ blockade. Additionally, while our early genetic investigations indicate CAL-101 treated T cells have a unique phenotype compared to vehicle or AKTi treated T cells, further genetic and epigenetic analysis will better elucidate if this phenomenon is due to preservation of a naïve-like phenotype, or true induction of a stem memory T cell phenotype.

Among the class 1A catalytic subunits of PI3K, P110δ is expressed mainly in the hematopoietic lineage (Vanhaesebroeck et al., 1997) while P110α and β are thought to play a minimal role (Ahmad et al., 2017; Okkenhaug, 2013; So et al., 2013). Yet, while CAL-101 is highly specific for p110δ we are cautious to attribute its potentiating effect on T cell memory solely to P110δ blockade, since CAL-101 also effects the class II, III and IV PI3Kinases when used at doses similar to what we used to treat our T cells (Lannutti et al., 2011). Future experiments with P110δ knockout mice with or without CAL-101 treatment will further elucidate the way in which CAL-101 primes T cells to exert more powerful antitumor immunity.

Inducing a durable memory phenotype through reversible pharmaceutical manipulation of PI3Kδ is an attractive way of generating potent memory T cells. Yet, while CAL-101 T cells exerted a longer antitumor response than vehicle and AKTi treated T cells, they eventually lost control of the tumor in most mice (Figure 2B). We found that as early as one week after transfer, these cells were expressing similar levels of PD-1 and KLRG-1 compared to vehicle and AKTi treated T cells (Figure 2A). Thus, while PI3Kδ blockade induces a strong memory phenotype including high Tcf7 expression, it does not appear to protect the cells from inhibitory signaling and exhaustion. We posit that a combination of PD-1 checkpoint blockade therapy may prevent the reestablishment of immune tolerance and subsequent tumor progression observed following treatment with CAL-101 T cells allowing for continued tumor control. Additionally, since CAL-101’s direct cytotoxic effects on cancers (Meadows et al., 2012) as well as its ability to break immune tolerance without affecting effector T cell viability (Ahmad et al., 2017; Ali et al., 2014) are well known, systemic administration of CAL-101 may synergize well with current checkpoint blockade or vaccine strategies. Nonetheless, whatever the route of application may be, we propose the use of CAL-101 as a viable and potent method for enhancing immune-based therapies for cancer to improve patient responses and survival.

## EXPERIMENTAL PROCEDURES

### Mice and cell lines

C57BL/6J (B6), pmel-1 TCR transgenic mice and NOD/scid/gamma chain knock out (NSG) mice were purchased from Jackson Laboratories. Mice were housed and experiments conducted in accordance with MUSC’s Institutional Animal Care and Use Committee’s (IACUC) procedures. B16F10 (H-2b) melanoma tumor, gift of the Nicholas Restifo lab at NCI surgery branch and M108 xenograft tumors were a gift from the June lab at the University of Pennsylvania.

### T cell cultures

<u>Pmel-1</u> CD8^+^ T cells were prepared from whole splenocytes were activated using 1μM hgp100 peptide + 100 IU rhIL-2/mL. Starting 3 hours after initial activation, cells were treated with either DMSO vehicle, AKT inhibitor VIII (AKTi) (Calbiochem), or CAL-101 (Selleckchem) at indicated doses. Cells were supplemented with culture media containing 100 IU rhIL-2/mL and vehicle or drug when expanded. <u>Human Normal Donor Peripheral T cells:</u> Polyclonal CD3^+^ T cells were activated using CD3/CD28 beads (Gibco) under 100 IU rhIL-2/mL DMSO vehicle, 1μM AKTi or 10μM CAL-101 and engineered with an anti-mesothelin 4-1BBξ chimeric antigen receptor (CAR, gift from the June lab). <u>Tumor Infiltrating Lymphocytes:</u> Tumor infiltrating lymphocytes (TILs) were derived from non-small cell lung cancer tumor samples from two patient donors provided by Dr. John Wrangle and Dr. Mark Rubinstein. Tumors were minsed into 1-3mm pieces, and transferred to wells of a 24-well plate containing CM with 6,000 IU/mL rrhIL-2 (TIL media). TIL were given fresh TIL media or split when confluent every 2-3 days for up to 5 weeks. On week 3, half of split wells were given CAL-101 (10 μM), while the original wells were treated with vehicle for the duration of the experiment.

### Adoptive cell therapy

<u>B6 mice</u> received 4.5e^5^ B16F10 cells subcutaneously 5-8 days before ACT. One day before therapy, mice underwent nonmyeloablative 5 Gy total body irradiation. CD8^+^ T cells were *in vitro* activated with feeder cells and peptide 12 hours before transfer via tail vein infusion. Unless otherwise indicated, IL-2 complex (see supplemental methods) was administered via intraperitoneal injections on days 0, 2, and 4 of treatment. In the antibody neutralization experiment, 200μg of either IL-7 neutralizing antibody (clone M25) or IgG2b isotype (clone MPC-11) (BioXCell) were administered by intraperitoneal injection on days 0, 3, 5, 8, 12, and 17 of treatment. <u>NSG mice</u> received 6e^6^ M108 suspended in matrigel subcutaneously 51 days prior to adoptive therapy. On day of treatment, mice received 3.5-4e^5^ CAR T cells via tail vein injection.

### In vivo Cytotoxicity Assay

Equal ratios of CTV (Thermo Fisher Scientific) labelled hgp100 and OTII-pulsed B6 splenocytes (0.5 and 5μM CTV label, respectively) were mixed at equal ratio then injected intravenously into mice previously treated with T cells or untreated control mice. After 4 hours, spleens were harvested from mice and labelled splenocyte frequencies assessed via flow cytometry and reported as % specific lysis using the following equation: % specific lysis = (1-(ratio of no T cell control mice) / (ratio of ACT mice)) x 100, where ratio = % OTII / % hgp100).

### Statistics

Kaplan-Meier survival curves were using a log rank test. Statistical comparisons between groups was performed via a student’s t-test for two groups, or a one-way ANOVA followed by multiple comparisons of group means (3+ groups). A p-value of <0.05 was considered significant. All statistics reported as mean ± SEM. Statistical analysis of differential gene expression was performed using Bonferroni-adjusted p-values to account for multiple hypothesis testing and a cut-off at 0.05 for the adjusted values to assign significance of the differential expression across conditions.

**Graphical Abstract.**
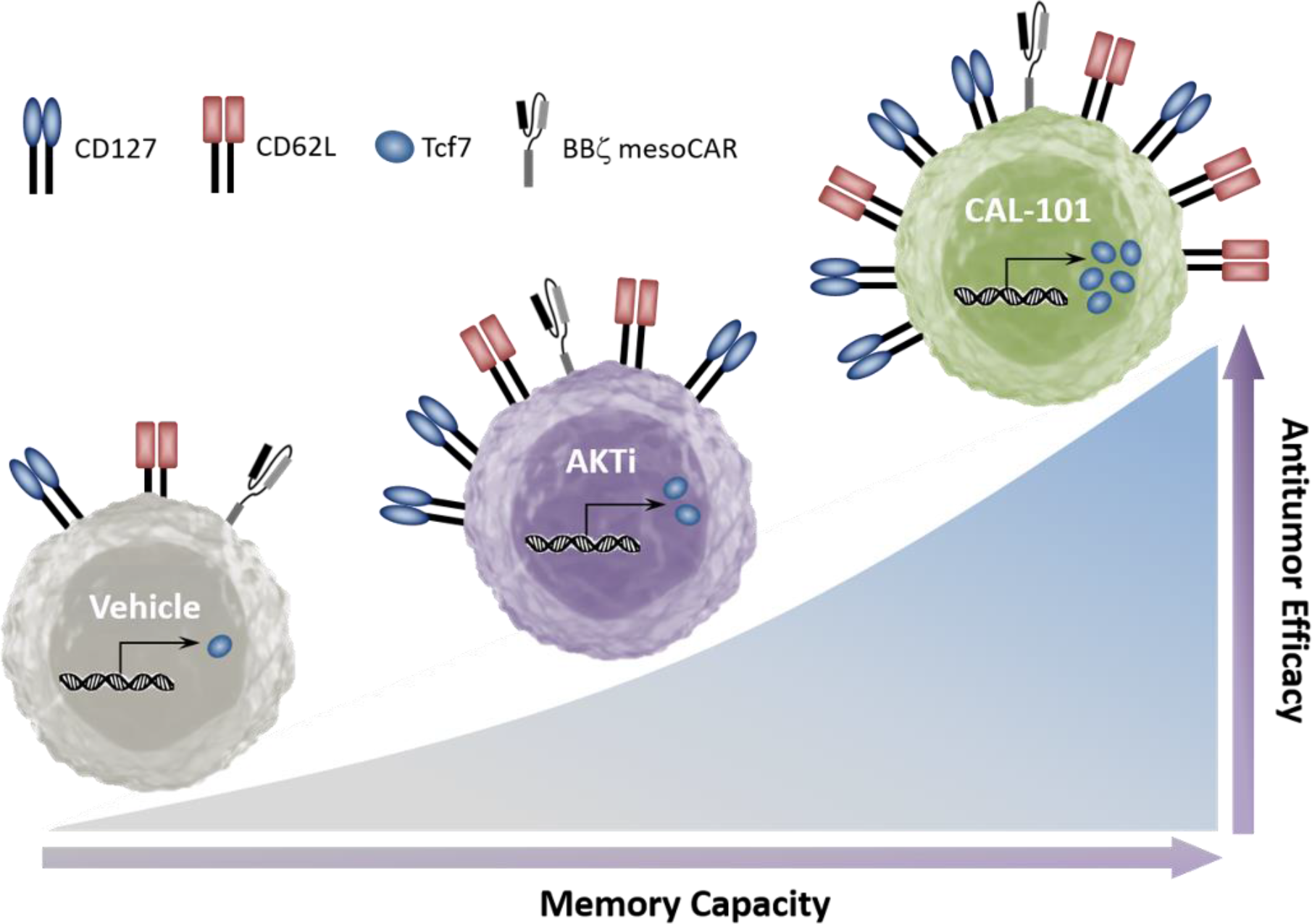
CAL-101 induces a strong memory phenotype and powerful antitumor capacity in CAR T cells. Graphical comparison of the memory capacity and antitumor efficacy of tumor-reactive T cells cultured in vehicle, AKT inhibitor (AKTi), or PI3Kδ inhibitor (CAL-101).

## AUTHOR CONTRIBUTIONS

Conceptualization, J.S.B., K.M. and C.M.P.; Methodology, J.S.B., K.M. and C.M.P.; Investigation, J.S.B., M.H.N., M.M.W. and A.S.S.; Formal Analysis, J.S.B., B.A.A. and J.H.; Writing – Original Draft, J.S.B. and C.M.P.; Writing – Review and Editing, J.S.B., K.M., M.H.N., B.A.A., M.M.W., A.S.S., S.R.B., L.N., J.H. and C.M.P.; Visualization, J.S.B., S.R.B. and C.M.P.; Funding Acquisition and Resources, J.S.B., M.H.N., S.R.B. and C.M.P.; Supervision and Project Administration, C.M.P.

## ACKNOWLEDGEMENTS

We thank Drs. John Wrangle and Mark Rubenstein for their collaboration in the experiments with human TIL. We further acknowledge Thomas Z. Benton for experimental support and Medgenome Inc. for mRNA transcriptome sequencing. This research was supported in part by the Hollings Cell Evaluation & Therapy, and Hollings Biostatistics Shared Resources, Hollings Cancer Center, Medical University of South Carolina (P30 CA138313).

This work was supported by NIH fellowship grant F30 CA200272 and NIH training grant T32 GM008716 to J.S. Bowers, a Jeane B. Kempner Foundation grant and ACS Postdoctoral fellowship (122704-PF-13-084-01-LIB) grant to M.H. Nelson, an NIH fellowship grant F31 CA192787 to S.R. Bailey, and NCI R01 grants R01 CA175061 and RO1 CA208514, KL2 South Carolina Clinical & Translational Research grant UL1 TR000062, ACS-IRG grant 016623-004, and MUSC Start-up funds to C.M. Paulos. The authors declare no conflict of interest.

